# An abundant future for quagga mussels in deep European lakes

**DOI:** 10.1101/2023.05.31.543086

**Authors:** Benjamin M. Kraemer, Salomé Boudet, Lyubov E. Burlakova, Linda Haltiner, Bas W. Ibelings, Alexander Y. Karatayev, Vadim Karatayev, Silvan Rossbacher, Raphael Stöckli, Dietmar Straile, Spaak Piet

## Abstract

Quagga mussels have expanded their range across the northern hemisphere in recent decades owing to their dispersal abilities, prolific reproduction rates, and broad ecological tolerances. Their remarkable capacity to filter particulates from the water column has had profound effects on inland aquatic ecosystems. In the North American Great Lakes, quagga mussel populations have increased inexorably since the late 1980’s, but it remains unclear whether quagga mussels will follow a similar trajectory in Europe where they have appeared more recently. Here we apply knowledge from a 33-year quagga population monitoring effort in the North American lakes to predict future quagga populations in deep European lakes, where quaggas are quickly becoming a conspicuous part of the underwater landscape. We predict that quagga mussel biomass in Lakes Biel, Constance, and Geneva may increase by a factor of 9 – 20 by 2045. Like in North America, this increase may be characterized by a shift to larger individuals and deeper depths as the population matures. If realized, this rapid expansion of quagga mussels would likely drive the largest aquatic ecosystem change in deep European lakes since the eutrophication period of the mid-20^th^ century.

## Introduction

The range of the quagga mussel (*Dreissena rostriformis bugensis* (Andrusov)) has rapidly expanded in the Northern Hemisphere since the 1980’s due to human activities that provided previously unavailable means of spread (Karatayev and Burlakova 2022b, Matthews *et al* 2014, Bij de Vaate *et al* 2013). After arriving in a new location, their spread is amplified by the downstream transport of their planktonic, free-swimming larvae (Karatayev and Burlakova 2022b, Matthews *et al* 2014). Quagga mussels settle on various substrates and tolerate a wide range of temperatures, salinity levels, and nutrient concentrations, allowing them to thrive in many different waterbodies. The ultimate success of quagga mussels after they appear in a waterbody is largely governed by the waterbody’s depth--lakes of similar maximum depth also have similar total quagga biomass, mussel sizes, and depth distributions (Karatayev and Burlakova 2022a, Karatayev *et al* 2021). This suggests that documented long-term quagga population dynamics from deep lakes in one part of the world could be used to estimate the likely quagga population trajectories in other lakes of similar depth.

But waterbody depth is not the only potential factor influencing quagga mussels. Other factors such as temperature, oxygen conditions, food availability, substrate availability, predation, and competition may also play a role. Thus, differences across a range of factors could preclude the use of a documented long-term quagga population dynamic in one lake to predict potential future dynamics in another. Here we apply knowledge from a 33-year quagga population monitoring effort in the North American Great Lakes to assess the probability that deep lakes in Europe will experience similar patterns based on comparisons of (1) their total biomass, mean individual mass, and biomass depth distributions early on post quagga detection, and (2) their climatological, morphometric, sedimentological, and biological characteristics. Depending on the results of those comparisons, we also project potential future quagga populations in deep lakes in Europe, where quagga populations have appeared more recently but are quickly expanding in size (Haltiner *et al* 2022).

## Method

### Study Areas

Quagga mussel population dynamics were studied in four lakes in North America (Erie, Huron, Michigan, Ontario) and three lakes in Europe (Biel, Constance, Geneva) (Fig 1). Quagga mussels were first detected in Lake Erie (1989) followed by Ontario (1990), Michigan/Huron (1997), Geneva (2015), Constance (2016), and Biel (2019) (Karatayev *et al* 2021, Haltiner *et al* 2022). These seven lakes represent a range of lake climate (average annual watershed precipitation: 829 – 1493 mm year^-1^; average watershed annual air temperature: 3.5 -7.1 °C), sedimentological (clay: 10-18 %; silt: 35 – 38 %; sand: 43 – 54 %), and morphometric characteristics (surface area: 38 – 59399 km^2^; mean depth: 16 – 156 m; maximum depth: 64 – 310 m (Lehner *et al* 2022)). The European lakes largely fall within the range of climatic, sedimentological, and morphometric characteristics of the North American lakes except in terms of their precipitation (European lake watersheds are wetter), sediments (European lakes have more clay and less sand), mean depths (Geneva and Constance are deeper), surface areas (European lakes are smaller) and watershed population densities (European lake watersheds have smaller total human populations but higher population densities) (Lehner *et al* 2022). The European lakes are also slightly warmer on average, and, unlike the North American lakes, have not experienced substantial ice cover since the end of the 20th century. The European lakes typically mix over a short period in February and March, whereas the North American lakes typically mix twice a year in autumn and spring. Due to the larger fetch, the mixing depths are deeper in the Great Lakes compared to the European lakes. The seven study lakes are biologically similar in the lack of major interspecific competitors for quagga mussels. Zebra mussels were present in all seven lakes prior to the detection of quaggas, but were largely displaced by them shortly after being detected (Haltiner *et al* 2022, Karatayev *et al* 2021, Matthews *et al* 2014). A variety of potential fish and bird predators inhabit all seven lakes, and the planktonic life stages of quagga mussels may be preyed on by fish and planktivorous microbiota. However, top-down limitation is likely restricted to the upper water column and is not considered an important factor influencing whole-lake quagga populations (Karatayev and Burlakova 2022b).

**Figure 1.**
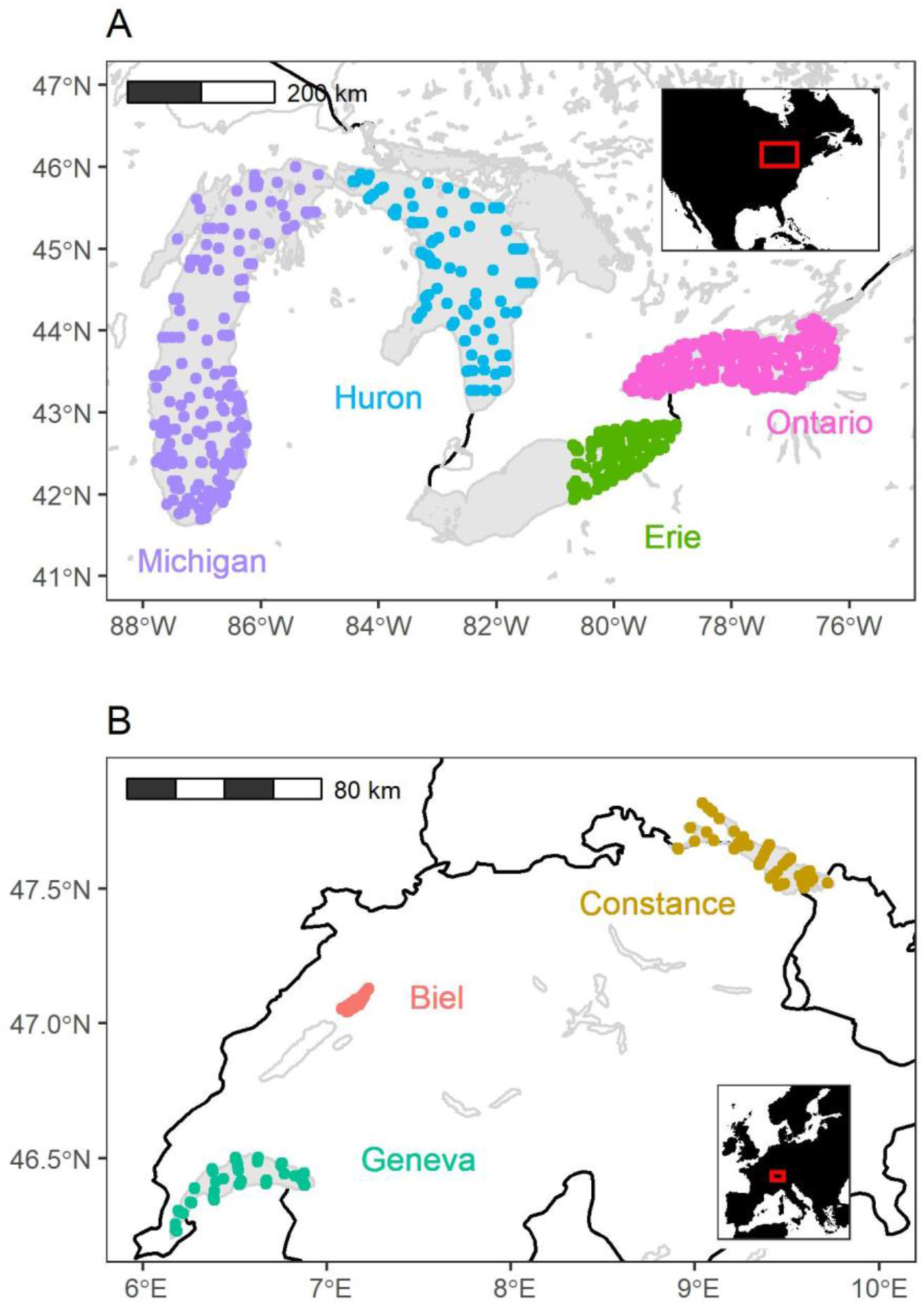
Map of the 1357 study sites showing the locations of quagga sampling efforts across 7 large lakes. Each circle represents a study site and the color of the circle represents the lake. Most study sites were sampled more than once except Biel and Geneva. Within each lake, the study sites are distributed across a large geographic area and depth range. Lake boundary map data come from the HydroLAKES database v1.0 which is licensed under a Creative Commons Attribution (CC-BY) 4.0 International License. (https://www.hydrosheds.org/products/hydrolakes).

Available data from the central and western basin of Erie were excluded from our analyses because they have distinct features that affect quagga mussel population dynamics (Karatayev *et al* 2021). The central basin of Lake Erie was excluded because regular hypoxia develops by the end of the growing season at depths > 20 m which restricts quagga populations to shallow areas (Karatayev *et al* 2018a). The western basin of Lake Erie was excluded because it is too shallow to be comparable to the other deep lakes in the analysis (Karatayev *et al* 2021). The shallow Saginaw Bay was also excluded from the greater Lake Huron as it has well documented differences in its ecology and human influence which affect quagga mussel population dynamics (Kraemer *et al* 2022, Karatayev *et al* 2021).

### Sampling

Quagga mussel samples were collected usually in triplicate at 1357 sites distributed across all lakes using a combination of grab sampling (PONAR or Ekman) and videography (Benthic Imaging System, BIS). The number of sampling points and sampling years varied across lakes (Biel: 29 sites and 1 year, Constance: 103 sites and 2 years, Geneva: 69 and 1 year, Huron: 171 points and 5 years, Michigan: 430 sites and 6 years, Ontario: 373 sites and 9 years, Erie eastern basin: 182 sites and 6 years). For the grab samples, all individuals retained following elutriation with a 1 mm (European lakes) or 0.5 mm (North American lakes) sieves were counted and data were reported as densities (individuals m^-2^). To ensure comparability across sites with different sieve sizes, all individuals smaller than 5 mm were excluded from our analysis. Quagga samples were collected using either regular (0.052m^2^) or petite PONAR (0.0483 m2) or Ekman samplers. All mussels were preserved in 10% formalin (North America) or frozen at -20°C (Europe) on the same day and later measured and counted in the laboratory. In all cases, mussels were further processed following the NOAA Technical Memorandum GLERL-164 (Nalepa *et al* 2014) and Standard Operating Procedure for Benthic Invertebrate Laboratory Analysis (US EPA 2015). For videographic samples, the percent coverage and number of individuals were obtained from bottom images recorded with BIS using Adobe Photoshop (for details see Karatayev *et al* 2022). To fully cover the time sequence since quagga detection, author-collected samples were supplemented with data from published literature for the North American lakes. Detailed sampling protocols for grab samples and videography can be found in the primary papers (Karatayev *et al* 2014, 2018a, 2018b, Nalepa *et al* 2014, Karatayev *et al* 2021, Nalepa *et al* 2018, 2020).

Ash-free dry weight (AFDW)--the biomass lost after oxidation– was used to compare quagga biomass across sites in a standard way which eliminates the variability introduced by the inorganic component of the sample, which can vary greatly depending on the type of sample and the conditions under which it was collected. The quagga mussels of each sample were separated from other species and dried for 48h at 60°C to obtain dry weight. Then the mussels, including their shells, were ashed at 550° for 2 hours and weighed. AFDW was calculated as the difference between dry and ash weight. In Lake Constance only 65 out of 103 sites included PONAR samples, so BIS density and percent coverage data were converted to a PONAR-equivalent AFDW estimate using statistical relationships captured with a General Additive Model (adjusted R^2^ = 0.90, p-value = <0.001, n = 1444). While quagga densities are available for all lakewide surveys conducted in the North American lakes, mussel weight was not always measured directly. In Lake Ontario, quagga were not weighed from 1990 to 2003, but weight was estimated using a constant conversion factor per mussel following Birkett et al. (2015) with some adjustments (Karatayev *et al* 2022). In Lake Huron, *Dreissena* biomass was not recorded from 2000 to 2007 and was estimated using an average of 5.19 mg AFDW per mussel, as calculated from 2012 samples (reviewed in Karatayev *et al* 2021). For consistency across studies, all historic and current data on quagga biomass were converted into AFDW in the same units (g m^2^) (Karatayev *et al* 2021) using standard equations (Glyshaw *et al* 2015, Nalepa *et al* 2014, Karatayev *et al* 2021, 2022).

### Modeling

We modeled quagga AFDW variation and mean individual weights (AFDW ind^-1^) as a function of the lake name, time in years since quaggas were detected, and the site-specific sampling depth in a 3-step model chain (Lin *et al* 2023). In the model chain, three models were fit in order of increasing complexity; first a linear model (LM), followed by a general additive model (GAM) followed by a boosted regression tree (BRT) where the residuals from the simpler model are used as the response variable in the next model so that, with each model, more of the variation is explained. The model chain included a combination of models to effectively balance the well-known tradeoffs between these statistical approaches. The summed predictions of all three models were used to generate the full set of bias-corrected predicted values (Figures 2-4) following previous work (Kraemer *et al* 2022). LMs and GAMs are known to be better at extrapolation and interpolation (Chow and Lin 1971). By fitting the LM and GAM first, we ensured that the extrapolative/interpolative component of the model performs as well as possible. But LM and GAM don’t fully capture the interactions between predictor variables, so we used BRTs as the final model in the stack to account for these complexities. When summed, the multi-model chain results in a “best of all worlds” scenario where the superior extrapolation performance of LM and GAM are combined with the superior capacity of BRTs to capture complexity in the predictor-response relationships.

**Figure 2.**
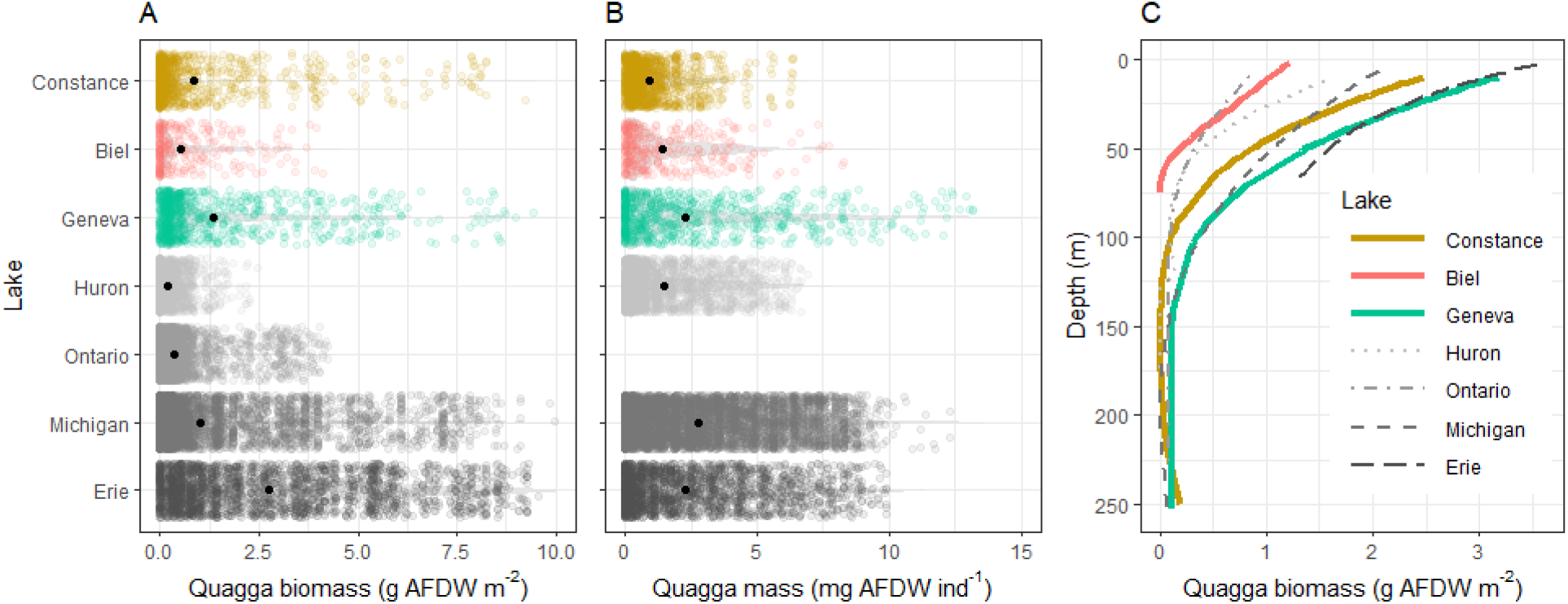
Modeled quagga total biomass (A), mean individual mass (B), and total biomass depth distributions (C) in deep European lakes follow similar patterns in deep North American lakes. Modeled quagga mussel biomass values represent those at 5 years after quagga mussels were detected to ensure comparability across lakes. Size data for Lake Ontario do not appear in panel B because it does not have sufficient size data needed to calculate quagga mass (mg AFDW ind^-1^).

**Figure 3.**
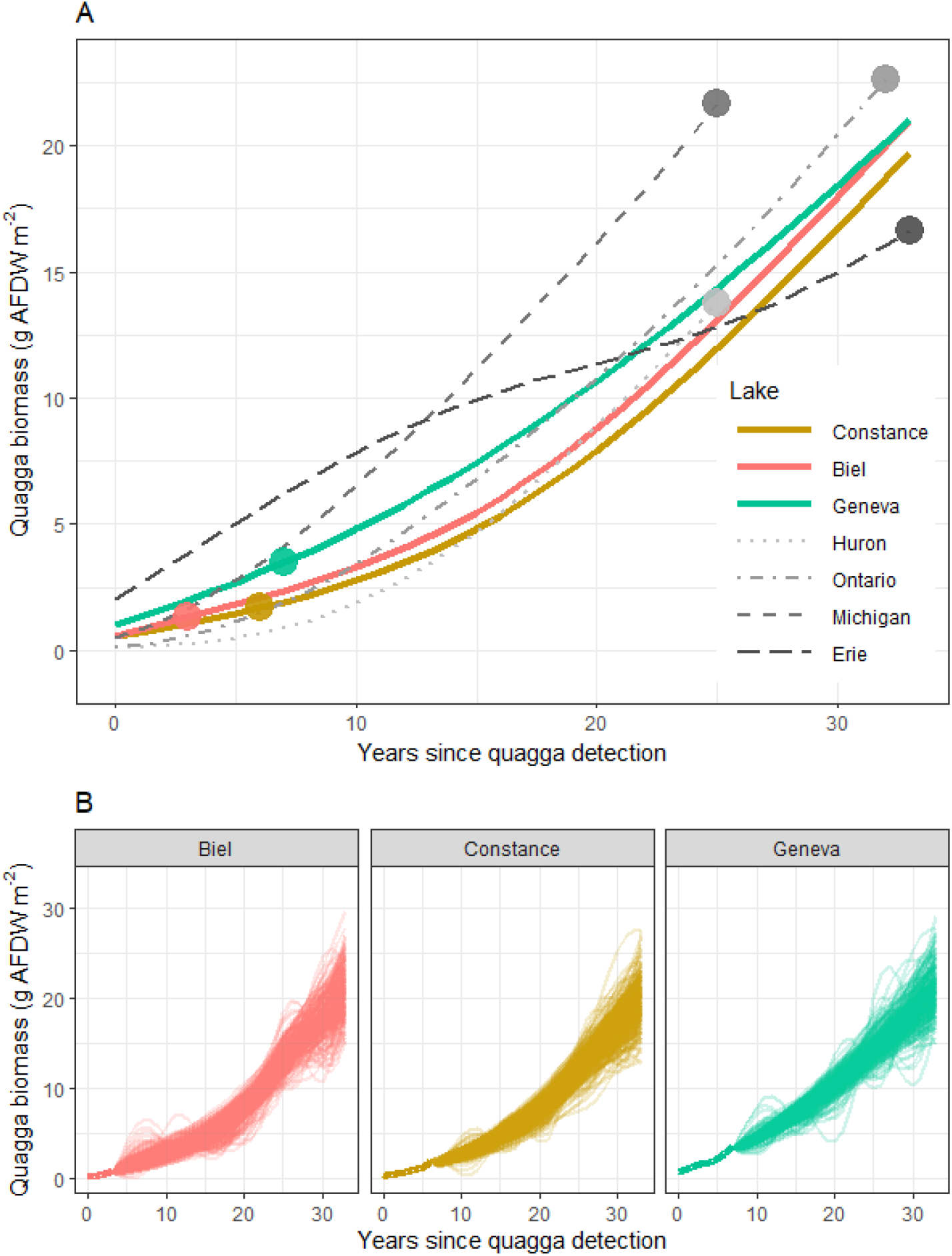
Predicted biomass of quagga mussels in each lake over a 33-year period suggest that Quagga mussel biomass will increase in deep European lakes. In panel A, each line represents the mean predicted biomass across all sites within each lake. The dot on each line represents the time since quagga detection specific to each lake in the year 2022. Panel B shows a range of possible outcomes (n = 1000) specifically for Lakes Biel, Constance, and Geneva after accounting for uncertainty due to random model errors using bootstrapped error propagation. The figure demonstrates that the biomass of quagga mussels is likely to increase in the future with modest differences among lakes in the magnitude of this increase. Lake basin colors and line types are consistent in all panels to aid visual comparison.

**Figure 4.**
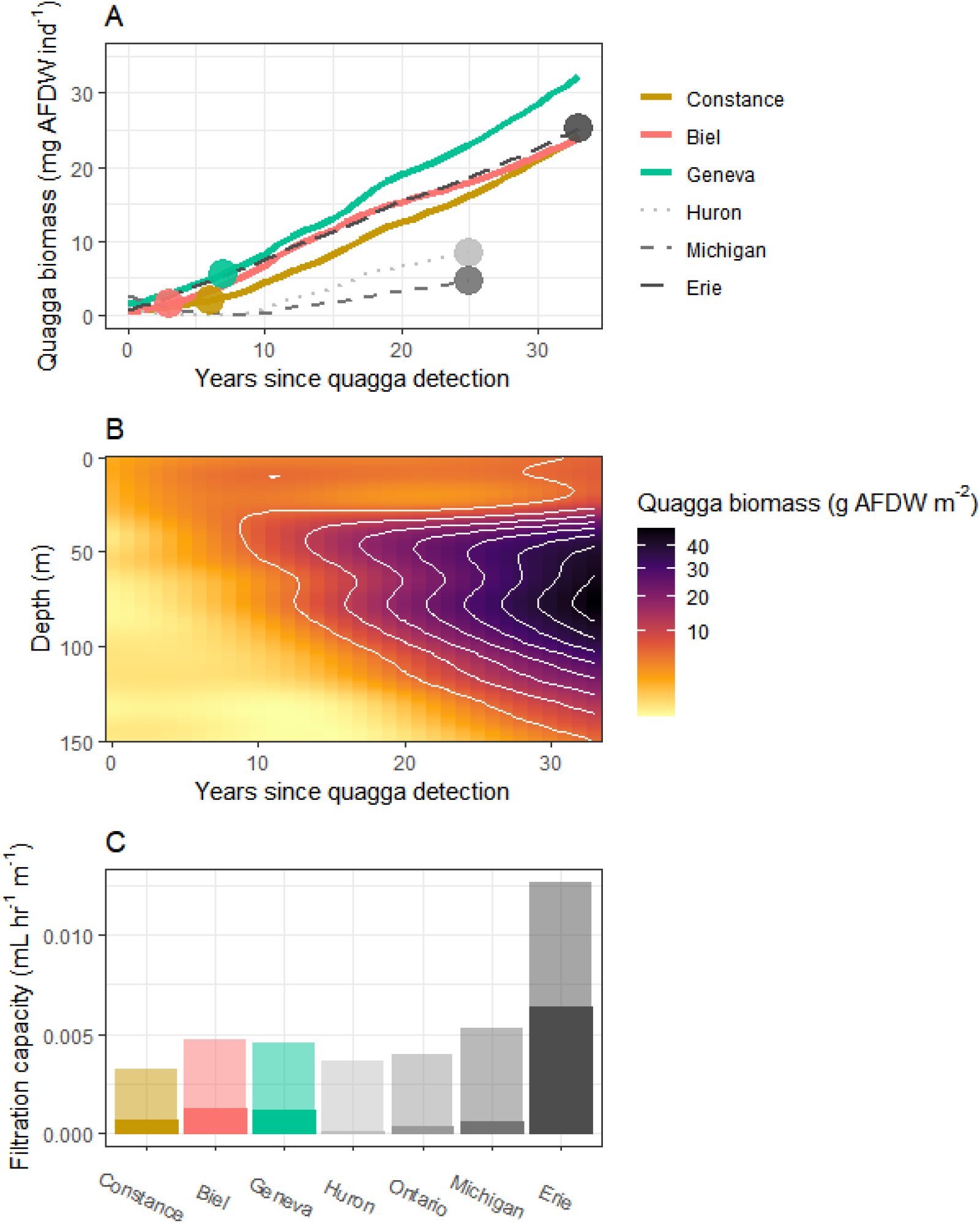
Over time, quagga mussels may become larger (A), found deeper (B), and with higher capacity to filter the water column (C). In panel A, each line represents the mean predicted individual biomass across all sites within each lake. Panel B shows the predicted change in quagga biomass across depth for Constance (Geneva and Biel show similar patterns). Panel C shows the modeled increase in filtration capacity between year 5 (solid bar) and year 30 (partially transparent bar) post quagga mussel detection.

The performance of the model chain was evaluated using cumulative adjusted R^2^ and median absolute deviation in cross validation with a 50-50 split between test and training datasets. Test and training datasets were randomly selected with 1000 repetitions and stratified by lake so that each lake’s data were split 50-50. To estimate the uncertainty in the future projected quagga biomass, we added randomly sampled values from the error distribution in the test datasets to the future modeled values of quagga biomass using a bootstrapped error propagation approach with 1000 repetitions. The model chain performed well when predicting quagga mussel biomass and mean individual mass with each successive model in the model chain explaining more of the variation in the response. In the model chain predicting variation in biomass (g AFDW m^-2^) across depth, lakes, and years since quagga detection, the cumulative adjusted R^2^ increased from 0.36 to 0.43 to 0.56 with the addition of each model in the stack. In the model chain predicting variation in mean individual mass (mg AFDW ind^-1^), the cumulative adjusted R^2^ increased from 0.38 to 0.47 to 0.51. The model chain also performed well in cross validation as the mean Spearman rank correlation between predicted and observed values in the test datasets were similar to the training datasets (0.60 versus 0.53 for biomass and 0.52 versus 0.43 for mean individual mass) with a median absolute deviation between predicted and observed values of 1.3 g AFDW m^-2^ and 1.9 mg AFDW ind^-1^ in the test datasets.

In an ideal case, raw data from each lake at a consistent time since detection would be used to compare early quagga population dynamics in each of the 7 lakes. However, quagga mussel sampling was made irregularly along the time sequence following detection, making it difficult to compare raw data at the same time since detection for all lakes. Instead, we used summed model output at year 5 since detection for each lake so that we could control for modest differences across lakes in the timing of their initial sampling events.

To determine the potential ecological impact of quagga mussels, we calculated the intensity of their filter feeding on phytoplankton during the spring and fall periods when the lakes are below 10 ॰C. We used modeled estimates of quagga mussel biomass and observed site-specific depths, along with moderate estimates of quagga mussel growth (0.06 g day^-1^) (Rowe *et al* 2017) and clearance rates (10 mL water mg AFDW^-1^ h^-1^) (Vanderploeg *et al* 2010) to calculate the fraction of the water column cleared per day by quagga filter feeding at each site. Our calculations were based on conditions typically observed during seasonal mixing where lakewide temperature differences are less than 10 °C, with moderate food quality, as reported in previous studies (Rowe *et al* 2017, Vanderploeg *et al* 2010, Karatayev *et al* 2021). These seasonal conditions were chosen because quagga filtration can have an especially potent effect on water clarity and productivity at that time of year (Karatayev *et al* 2021).

We used the statistical software R (R Core Team 2022) for statistical computing and data analysis. We used several R packages including the ‘data.table’ package for data manipulation (Dowle and Srinivasan 2021), the ‘ggplot2’ package for data visualization (Wickham *et al* 2016), the ‘mgcv’ package for general additive modeling (Wood and Wood 2015), and the ‘dismo’ and ‘gbm’ packages (Hijmans *et al* 2017, Greenwell *et al* 2019, Elith *et al* 2008) for boosted regression tree modeling.

## Results

Despite the numerous differences between the climate, sediments, and morphometry, early population dynamics suggest that European lakes have thus far followed a similar path to that of the North American lakes (Fig 2). The quagga total AFDW in the European lakes fell within the range of the North American lakes and was most similar to lakes Huron, Michigan, and Ontario (Fig 2a). European lakes also had similar mean individual quagga mass (mg AFDW individual^-1^; Fig 2b). The quagga AFDW depth distribution in the North American lakes peaked above 50 m and decreased substantially with depth (Fig 2c). The depth distribution of quagga biomass in Biel was most similar to Ontario/Huron, Constance was most similar to Michigan, and Geneva was most similar to Erie (Fig 2c). These similarities (Fig 2a-c) suggest that quagga population dynamics in the European lakes have followed those of the North American lakes, providing justification for further analyses projecting the potential future of quaggas in lakes Biel, Constance, and Geneva.

If quagga populations continue on a course similar to the North American lakes, quagga biomass in Europe is expected to increase in the coming decades (Fig 3). We find that quagga mussel biomass in Biel, Constance, and Geneva may increase by a factor of 9 – 20 by 2045. The projected biomass at year 30 post quagga detection were similar in Lake Geneva (18.4 g m^-2^, 95% CI: 14.4 - 22.4 g m^-2^) Biel (17.9 g m^-2^, 95% CI: 12.9 – 22.3 g m^-2^) and Constance (16.8 g m^-2^, 95% CI: 12.7 - 20.9 g m^-2^). As in the case of the early population dynamics post quagga detection (Fig 2), the European lakes are most similar to lakes Ontario and Huron in their projected increases in quagga biomass (Fig 3a).

Like in the North American lakes, the projected increase in quagga biomass will likely be characterized by a shift to larger individuals, deeper depths, and higher filtration capacity as the quagga population matures (Fig 4). The average quagga mass for the European lakes is projected to increase by a factor of 8.2 from 2.9 mg AFDW ind^-1^ at 5 years post quagga detection to 23.7 mg AFDW ind^-1^ at 30 years post quagga detection. Over the same time interval, the depth with the highest mussel biomass (g AFDW m^-2^) is projected to shift deeper from 14 m to 78 m on average across the European lakes. As a first approximation, these changes are expected to increase the capacity of the quagga population to filter the water column during seasonal mixing by 370 - 510 %, potentially resulting in strong effects on the water column phytoplankton biomass, suspended sediment, and overall water clarity. Taken together, these changes may strongly affect the ecosystem functioning of European lakes in the coming decades.

## Discussion

Based on the comparisons with invaded lakes in North America, we estimate that the cumulative quagga biomass across lakes Geneva, Biel, and Constance will increase to approximately 18 million kg of AFDW over the course of the 3 decades since they were detected (sum of mean lakewide quagga AFDW per unit area at year 30 post quagga detection * surface area of each lake) -- a total weight approximately equivalent to 3,150 fully-grown male African elephants. If realized, this projected expansion of quagga mussels would drive the largest aquatic ecosystem change in deep European lakes since the eutrophication period of the mid-20th century. But unlike the problem of eutrophication which had an apparent and feasible solution (i.e. reduce nutrient pollution), quagga mussels are a less tractable target for management *ex post facto* because their appearance in lakes is largely considered irreversible. Because of its irreversibility, quagga mussel populations are likely to be a flashpoint in lake management until they equilibrate with the local biological community which may not occur for decades. In the meantime, we encourage the development and implementation of a comprehensive management plan that takes into account the expected local effects of quagga on lake ecosystems and the benefits that people derive from those ecosystems (Burlakova *et al* 2022, Boltovskoy *et al* 2022). We also caution that the ultimate consequences of quagga mussels in Europe will depend on how they interact with other neobiota and other anthropogenic forces such as climate change.

The major ecosystem consequences of quagga mussels include increased water clarity, modified food webs, altered hydrodynamics, and changes to biogeochemical cycles (Hecky *et al* 2004, Karatayev *et al* 2021, Li *et al* 2021). Most of these changes arise from their efficient filter feeding which removes zooplankton, phytoplankton, suspended organic material, and sediment from the water column. By filtering these particulates from the water column, quaggas can dominate the uptake of lake nutrients, offsetting or even overshadowing the effects of external nutrient loading (Li *et al* 2021). Their filtration also allows light to penetrate deeper where it supports benthic primary production and may also stimulate benthic mats of toxic cyanobacteria (Francoeur *et al* 2015). Food webs are further altered by reducing the availability of food for planktivorous animals with cascading effects throughout the food web. But, quagga mussels themselves can be an important food source for some fish (Baer *et al* 2022a, 2022b, Karatayev and Burlakova 2022b, 2022a). For instance, in Lake Constance, benthic whitefish (*Coregonus macrophthalmus*), roach (*Rutilus rutilus*) and tench (*Tinca tinca*) are all capable of high levels of quagga consumption (Baer *et al* 2022a, 2022b). Waterfowl populations may also increase as they feed on quaggas that grow in shallow areas of lakes (Molloy *et al* 1997, Petrie and Knapton 1999).

The effects of quagga filter feeding has consequences for how people will interact with deep European lakes in the future. For instance, clearer water can be more inviting for swimming, boating, and scuba diving, although quaggas also foul beaches, boats, docks and piers. Increases in water clarity can increase the value of shoreline properties (Limburg *et al* 2010, Burlakova *et al* 2022). Water infrastructure including collecting stations for drinking water are typically placed around 60 meters depth in deep European lakes because the bivalves at the time of construction were restricted to the upper 40 m of water (Wacker and Von Elert 2003). According to our projections, quagga mussels will soon become abundant down to 100 meters or more, requiring millions of euros annually to cover damages to water intakes alone. Existing and future water infrastructure projects in the lakes must account for these additional costs.

By affecting food availability, changing habitats, and altering food webs, quagga mussels may also affect fishery productivity (Chiapella *et al* 2023, Lauber *et al* 2020). But the ultimate effect of quaggas on fisheries productivity in Europe will depend on how they interact with other ongoing environmental changes. For example, the re-oligotrophication of lakes in Europe has already caused changes in lake fisheries in recent decades (Eckmann *et al* 2006, Baer *et al* 2017), and these effects may be further compounded by quagga mussels. The three-spined stickleback (*Gasterosteus aculeatus*), first detected in the open water fish community of Lake Constance in 2013, and quickly became the dominant fish species (50% -70% of all fish individuals are stickleback), and may have already reduced Lake Constance fish catches (Baer and Brinker 2022). Quaggas may exacerbate or mitigate the effects of three-spined sticklebacks on pelagic fisheries, but at the moment, such anticipated effects are speculative. However, interactions between quagga and other neobiota may increase fishery productivity (Karatayev and Burlakova 2022b, 2022a, Madenjian *et al* 2015). For instance, Round gobies consume quagga mussels (Kornis *et al* 2012, Wilson *et al* 2006) and are in turn an important food source for commercially and recreationally valuable fish species including lake trout, burbot, yellow perch, smallmouth bass, lake sturgeon, and walleye (Karatayev and Burlakova 2022a). Round gobies are currently not present in the three European lakes included here, but may appear soon (Kalchhauser *et al* 2013) with a potential to limit quagga biomass in shallow areas and boost fishery productivity.

Quagga mussels may also interact with the anticipated effects of climate change on hydrodynamics. For instance, climate change is expected to reduce lake mixing leading to reductions in deep water oxygen (Jane *et al* 2021) that could limit quagga depth distributions or even their total populations as has occurred in the central basin of Lake Erie (Karatayev *et al* 2018a). Lake Geneva in particular is expected to have more widespread anoxia especially below 150 m (Schwefel *et al* 2016, Deyle *et al* 2022) which will limit quagga populations.

Episodic anoxia could affect quagga survival and reproduction at shallower depths as well. The capacity of quaggas to filter the water column would also depend on climate change’s anticipated effects on lake hydrodynamics. Periods of lake mixing are an important time when quaggas have potent effects on water column suspended material (Rowe *et al* 2015). Thus climate change-mediated reductions in lake mixing as predicted for the European lakes (Peeters *et al* 2007, Kraemer *et al* 2015) could reduce the capacity of quaggas to filter the entire water column during these critical seasonal periods. The consequences of higher temperatures for the mussel metabolism and growth are also unclear as higher temperatures typically lead to higher metabolic rates (Kraemer *et al* 2017), but quagga mussel metabolism and growth rates may be suppressed if not supported by food supply (Karatayev *et al* 2018b). Thus, the cumulative effects of climate change on quagga mussel populations and their ecosystem consequences are highly uncertain.

Although our models suggest that quagga population dynamics in deep North American and European lakes are similar in the first several years post quagga detection, we expect that, in the European lakes, that the future quagga biomass may be higher because they have higher chlorophyll-a (chl-a) concentrations, except for Lake Erie (Supplementary table 1). Higher average chl-a concentrations could make the European lakes more suitable for quagga and may cause higher quagga total biomass in the long run (Karatayev *et al* 2018b). This potential future boost to quaggas in deep European lakes may have the largest effects on Geneva and Constance whose substantial depth offers greater potential for quaggas to escape the effects of benthivorous birds and fish. However, the lower chl-a in the North American lakes may already be a consequence of quagga mussels rather than an indicator of their lower suitability for quagga. Nonetheless, to better constrain how regional differences in lake chl-a might affect future quagga populations, we encourage global investigations into the relationship between lake productivity, quagga populations, and their ecosystem impacts.

## Conclusions

An abundant near-future may lie ahead for quagga mussels in deep European lakes. The ultimate ecosystem consequences of quagga mussels in Europe will depend on their complex interactions with the local context and other anthropogenic drivers of change. But based on past dynamics in other lakes, quaggas in lakes Biel, Constance, and Geneva are likely to increase water clarity, damage important water infrastructure, alter fisheries productivity, and redirect nutrients and energy flow. In North American lakes, the strong ecosystem consequences of quagga mussels started to appear approximately 10 years after they were first detected (Karatayev *et al* 2022). Thus, it may still be too early to detect substantial changes in European lakes now, in less than 10 years after they first appeared. But, with new tools like remote sensing, high frequency monitoring platforms, and eDNA, managers and scientists are more equipped today to monitor and detect the ecological consequences of quagga mussels than ever before. Whether the ecological consequences of quagga mussels are harmful or beneficial for European society ultimately depends on the values of the various stakeholders in the region. We encourage careful consideration of both the costs and benefits of quagga mussels in the development of a comprehensive multinational management strategy (Sax *et al* 2022, Schlaepfer *et al* 2011). The window of opportunity to develop and implement such a plan is narrowing.

## Acknowledgements

This study was supported/co-financed by the grant “SeeWandel: Life in Lake Constance - the past, present and future” within the framework of the Interreg V programme “Alpenrhein-Bodensee-Hochrhein (Germany/Austria/Switzerland/Liechtenstein)” which funds are provided by the European Regional Development Fund as well as the Swiss Confederation and cantons. LEB was supported by the Prime Agreement Award GL 00E02259-2 from the U.S. EPA “Great Lakes Long-Term Biological Monitoring Program 2017-2022” (PI L. Rudstam, Co-PIs Burlakova and Karatayev). The funders had no role in study design, data collection and analysis, decision to publish, or preparation of the manuscript.

## Data and code availability statement

The data and code used in this study will be included as supplementary material accompanying this paper upon acceptance. Private links to the data and code will be shared with the target journal for the purpose of peer review.

## Ethics statement

This study involved the collection of live quagga mussels (*Dreissena rostriformis bugensis* (Andrusov)) from their natural habitat. The research procedures and handling of the mussels were conducted in accordance with the ethical guidelines and regulations governing animal research.

## Supplementary Material

**Supplementary Table 1:** Lake characteristics of the 7 lakes. Data are from the lake atlas database (Lehner et al 2022) and Kraemer et al (2022).

| Lake name | Michigan | Ontario | Huron | Erie | Constance | Geneva | Biel |
| --- | --- | --- | --- | --- | --- | --- | --- |
| Lake ID (from HydroLAKES) | 6 | 7 | 8 | 9 | 1243 | 1261 | 14023 |
| Country | United States of America | United States of America, Canada | United States of America, Canada | United States of America, Canada | Switzerland , Austria, Germany | Switzerland , France | Switzerland |
| Continent | North America | North America | North America | North America | Europe | Europe | Europe |
| Lake area (km<sup>2</sup>) | 57726.8 | 19347.4 | 59399.3 | 25767.8 | 522.02 | 571.63 | 37.59 |
| Shore length (km) | 2862.67 | 2609.93 | 8856.64 | 1935.52 | 284.79 | 177.97 | 46.79 |
| Shoreline development index | 3.36 | 5.29 | 10.25 | 3.4 | 3.52 | 2.1 | 2.15 |
| Total volume<br>(km<sup>3</sup>) | 4860000 | 164000<br>0 | 3550000 | 499000 | 49000 | 89000 | 617.78 |
| Impounded<br>volume (km<sup>3</sup>) | 0 | 29960 | 0 | 0 | 0 | 0 | 0 |
| Average<br>depth (m) | 84.2 | 84.8 | 59.8 | 19.4 | 93.9 | 155.7 | 16.4 |
| Maximum<br>depth (m) | 281 | 244 | 229 | 64 | 251 | 310 | 74 |
| Residence<br>time (days) | 29956 | 2450.5 | 4484.5 | 589.5 | 1626.2 | 3958.7 | 34 |
| Elevation<br>above sea<br>level (m) | 175 | 73 | 175 | 172 | 392 | 370 | 425 |
| Mean annual<br>watershed air<br>temperature<br>(°C) | 7.1 | 5.7 | 4.9 | 5.5 | 5.0 | 3.5 | 5.6 |
| Mean annual<br>watershed<br>precipitation<br>(mm year<sup>-1</sup>) | 830 | 850 | 829 | 842 | 1233 | 1493 | 1295 |
| Mean watershed soil clay fraction (%) | 13 | 11 | 10 | 11 | 16 | 16 | 18 |
| Mean watershed soil silt fraction (%) | 38 | 37 | 35 | 36 | 38 | 38 | 38 |
| Mean watershed soil sand fraction (%) | 47 | 50 | 54 | 49 | 44 | 45 | 43 |
| Shoreline population (1,000's) | 1855.47 | 1671.37 | 346.745 | 965.087 | 385.849 | 628.039 | 60.058 |
| Watershed population (1,000's) | 11126.1 | 37653.5 | 14826.7 | 27220.6 | 1655.72 | 1137.97 | 1151.46 |
| Shoreline human footprint (%) | 16.5 | 18.7 | 8.9 | 23.7 | 29.1 | 35.5 | 28.1 |
| Watershed<br>human<br>footprint (%) | 9 | 7.4 | 5.1 | 6.8 | 16.9 | 16.4 | 18 |
| Mean annual<br>surface chl-a<br>concentration<br>(µg L<sup>-1</sup>) (1997-2020) | 0.79 | 0.97 | 0.62 | 2.46 | 2.71 | 2.36 | 2.40 |

